# TERL: Classification of Transposable Elements by Convolutional Neural Networks

**DOI:** 10.1101/2020.03.25.000935

**Authors:** Murilo Horacio Pereira da Cruz, Douglas Silva Domingues, Priscila Tiemi Maeda Saito, Alexandre Rossi Paschoal, Pedro Henrique Bugatti

## Abstract

Transposable elements (TEs) are the most represented sequences occurring in eukaryotic genomes. They are capable of transpose and generate multiple copies of themselves throughout genomes. These sequences can produce a variety of effects on organisms, such as regulation of gene expression. There are several types of these elements, which are classified in a hierarchical way into classes, subclasses, orders and superfamilies. Few methods provide the classification of these sequences into deeper levels, such as superfamily level, which could provide useful and detailed information about these sequences. Most methods that classify TE sequences use handcrafted features such as k-mers and homology based search, which could be inefficient for classifying non-homologous sequences. Here we propose a pipeline, transposable elements representation learner (TERL), that use four preprocessing steps, a transformation of one-dimensional nucleic acid sequences into two-dimensional space data (i.e., image-like data of the sequences) and apply it to deep convolutional neural networks (CNNs). CNN is used to classify TE sequences because it is a very flexible classification method, given it can be easily retrained to classify different categories and any other DNA sequences. This classification method tries to learn the best representation of the input data to correctly classify it. CNNs can also be accelerated via GPUs to provide fast results. We have conducted six experiments to test the performance of TERL against other methods. Our approach obtained macro mean accuracies and F1-score of 96.4% and 85.8% for the superfamily sequences from RepBase and 95.7% and 91.5% for the order sequences from RepBase respectively. We have also obtained macro mean accuracies and F1-score of 95.0% and 70.6% for sequences from seven databases into superfamily level and 89.3% and 73.9% for the order level respectively. We surpassed accuracy, recall and specificity obtained by other methods on the experiment with the classification of order level sequences from seven databases and surpassed by far the time elapsed of any other method for all experiments. We also show a way to preprocess sequences and prepare train and test sets. Therefore, TERL can learn how to predict any hierarchical level of the TEs classification system, is on average 162 times and four orders of magnitude faster than TEclass and PASTEC respectively and on a real-world scenario obtained better accuracy, recall, and specificity than the other methods.

## Introduction

Transposable elements (TEs) are sequences present in genomes that can change location and produce multiple copies of themselves throughout the genome [9,26]. By doing so, these sequences are responsible for several effects, such as co-opt with the regulation of gene expression during grape development [11,21], regulate the classic wrinkled character in pea [5,8], and can increase the resistance to DDT insecticide of the *Drosophila melanogaster* [3,4].

There are 29 types of TEs, which can be classified in a hierarchical way into classes, subclasses, orders, and superfamilies, as proposed by Wicker et al. [26] (Table 1).

**Table 1.**
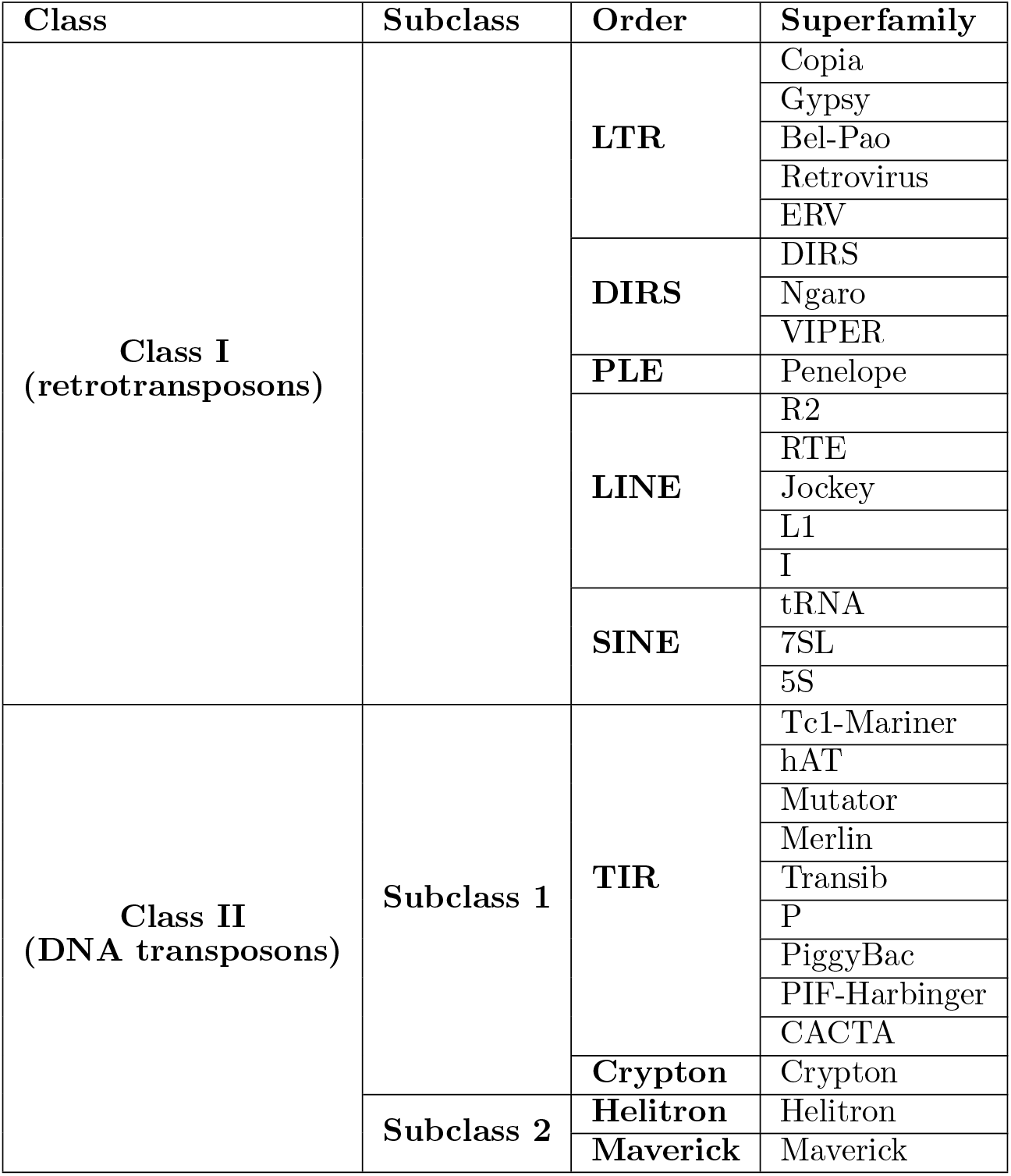
Classification of eukaryotic transposable elements [26]. Columns represent the hierarchical division from left to right.

Most methods that propose an automatic classification of TE sequences are based on manually pre-defined features, such as TEclass [1], PASTEC [14] and REPCLASS [10]. All these methods use homology based search as one of the features to classify TE sequences. The use of prior information requires databases with a huge amount of sequences from unrelated organisms, to provide a good search space.

### Contributions

In this paper, we present an efficient and flexible approach, transposable elements representation learner (TERL) to classify transposable elements and any other biological sequence. This work also presents the following contributions:

1. Sequence transformation to image-like data;
2. The use of deep convolutional neural networks as a feature extraction algorithm;
3. The use of deep neural networks as classifier;
4. Evaluation of coherence between different databases;
5. Creation and evaluation of data sets with sequences from different databases;
6. Generic classification to any other level and any biological sequence.

CNNs are representation learning and feature extraction algorithms [12,20,29], they do not require the use of any homology based search or any handcrafted feature and they are well applied to other genomic sequences prediction problems, as showed in [7,16,22,28].

In this paper, we propose the pipeline TERL to classify transposable elements sequences and any other biological sequence. The pipeline is composed by four preprocessing steps, a transformation from one-dimensional space data (i.e., nucleic acid sequences) into two-dimensional space data and a convolutional neural network classifier. We apply the pipeline to classify TE sequences into class, order and superfamily levels. The proposed pipeline does not require the use of any homology based search into known sequences databases nor the use of a series of models for each hierarchical level.

We also analyzed the performance obtained by TERL on a series of experiments and compare the results with the performances obtained by TEclass and PASTEC.

This paper is organized as follows. Section briefly presents some concepts of CNNs, the use of CNNs in genomics and related works. Section presents the proposed approach to classify TE sequences by using CNNs. Section presents the data sets, describes the experiments proposed to analyze the performance of TERL and shows the results obtained, including comparisons with other methods. Finally, on Section, we present our conclusions.

## Background

### Convolutional Neural Networks

CNN is a deep learning algorithm that can be applied to several tasks, like image classification, speech recognition and text classification [18]. It is also a representation learning algorithm, which can learn how to represent the data to perform some task by extracting features from it.

The architecture of CNN is inspired by the visual cortex of vertebrates, which is structured in a way that simple cells can recognize simple structures, like edges and lines in images, and based on what was recognized by these simple cells, other cells can recognize more intricate structures [12] [20].

The architecture is basically composed by an artificial neural network with interconnected units arranged in a specific way in multiple layers, which usually are: input, convolution, pooling and fully connected. The input layer is the normalized numerical representation of the raw data.

The convolution layer is composed by artificial neuron units connected to sliding window filters (i.e. feature maps) which apply convolution to small regions of its input data. These regions are defined based on the dimensions of the filter and the stride (i.e. amount of data to be skipped before the next convolution operation).

The convolution operation is defined by the dot product between the input layer data comprised within the region of the filter, *X*, and its weights, *W*. The sum between the dot product and a bias value is applied to an activation function, as Equation 1 shows. Usually the activation function used in a convolution layer is the non-linear function ReLU, which is defined by Equation 2.

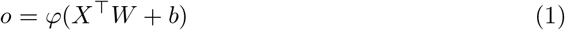

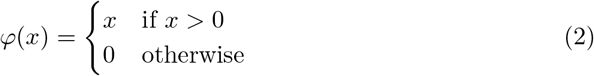

The application of convolution operations in small regions throughout the input data gives the CNN spatial connections property, which means that the network is able to learn how to recognize patterns independently of their amount and location in the input data. The number of trainable parameters is also reduced, which improves training efficiency.

The pooling layer usually applies a non-overlapping downsampling sliding window filter in its input data, which reduces the dimension of it by summarizing the values within each region. The two most used types of pooling layer are the max pooling, which extracts the maximum value within the region and the average pooling, which extracts the average of the region.

The downsampling applied by a pooling layer reduces even more the amount of trainable parameters as the next input layer data will be smaller and the pooling layer does not have any trainable parameter. This layer also provides local translation invariance as it summarizes the values of the regions to a single output value.

The fully connected layers basically compose a multilayer perceptron (MLP). Its input layer is the extracted features from the convolution and pooling layers. These layers are responsible to classify the input data.

The network updates its weights by minimizing some loss function, which calculates the distance between the outputs of the network and the desired outputs. This process allows the network to approximate its model to a desired model.

The method is trained by applying training samples in the network with their respective labels. The network calculates the outputs based on its parameters values (i.e. weights), and these parameters are updated by minimizing the loss function with some stochastic gradient descent based algorithm.

The performance of the trained model is tested by applying test samples in the network with their respective labels and checking its ability to predict the correct labels. The training set and the test set must not overlap to avoid overfitting and misleading performance results.

### CNNs in Genomics

CNNs are major applied to solve computer vision problems and other pattern recognition areas such as speech recognition. However, there are also studies that apply CNNs in biological problems [29].

In bioinformatics, CNNs can learn the regulatory code of the accessible genome, as discussed in [16]. This work developed an open source package, Basset, that use CNNs to learn the functional activity of DNA sequences. This method can learn the chromatin accessibility in cells and annotate mutations. Basset surpassed the predictive accuracy of the state-of-the-art method, showing how promise deep CNNs can be when applied to bioinformatics problems.

Deep CNNs also obtained very promising results on the prediction of DNA-protein binding, as showed in [28]. This work explores different architectures and how to choose the hyperparameters of the architecture. In the present study we show that adding convolution kernels to the network is important for motif-based tasks such TEs classification.

Identifying and modeling the properties and functions of DNA sequences is a challenging task. A hybrid convolution and bi-directional long short-term memory recurrent deep neural network framework, DanQ, is proposed in [22] to address this problem. This work shows how promising CNNs are as the method improved accuracy, recall, specificity and classification time compared to the results of other methods.

### Related Works

#### TEclass

TEclass [1] classifies TE sequences into DNA transposons (Class II) class and into Long Terminal Repeats (LTR), Long Interspaced Nuclear Elements (LINE) and Short Interspaced Nuclear Elements (SINE) orders.

The method consists of four binary classification steps for three sequence length ranges: 0-600, 601-1800, 1801-4000 bp; and three binary classification steps for sequences longer than 4000 bp. Each binary classification step consists of two Support Vector Machines (SVMs) models that perform classification using tetramers and pentamers (i.e. k-mer with k equals four and five). The classification steps follow a hierarchical order, in which each step can produce an output or proceed to the next step.

TEclass uses sequences from RepBase version 12.11 to build the SVM models and tests its performance with the new sequences added on RepBase version 13.06. TEclass obtained accuracies of 90.9%, 94.3%, 74.1% and 84.2% for Class II, LTR, LINE and SINE sequences, respectively.

#### PASTEC

PASTEC [14] is a method based on Hidden Markov Models (HMMs) profiles and classify TE sequences into two classes and 12 orders.

The features used to describe TE sequences are: sequence length, long terminal repeat (LTR) or terminal inverted repeat (TIR), Simple Sequence Repeats (SSRs), polyA tail and Open Reading Frame (ORF). The method also uses a homology search in RepBase Update with nucleotides and amino acids sequences and homology search in HMMs profiles of protein domains. The homology based searches are performed by blastx, tbastx and blastn.

Each of the homology searches can be turned off when classifying sequences, and thus the results can be affected, as we demonstrate on experiment 6 on Section. The choice of the auxiliary search files can also affect the results of the method since we show that PASTEC mostly depends on homology search.

According to [14], PASTEC obtained 63.7% and 51.3% of accuracy for the classification of sequences present in RepBase version 15.09 (that are not present in version 13.07) into class and order levels, respectively. This method also presents a misclassified rate of 2.9% and 10.9% for the classification of the same sequences into class and order levels, respectively. The method obtained lower misclassified rates compared to TEclass and REPCLASS.

### REPCLASS

REPCLASS [10] basically can classify TEs into every known category. The method consists of three basic modules that classify a given sequence independently. The class of a sequence is determined based on the consensus of three modules.

The first module, homology, consists of a homology search to known TEs in RepBase performed via tblastx. The second module, structure, consists of a search for structural elements in the query sequences and based on these structural evidences it classifies them. The third module, Target Site Duplication (TSD), consists of extracting individual copies from the target genome sequence and searching for TSDs in their flanking sequences.

REPCLASS obtained 26.1% and 66.9% of accuracy for the same data set used to evaluate PASTEC on the classification into class and order levels [14], and misclassified rates of 31.8% and 11.1%.

## Proposed Method

Here we propose an automatic classification of TE sequences by using TERL, which is composed of four preprocessing steps, a data transformation and convolutional neural networks. All the steps of the proposed pipeline are shown in Figure 1. In order to create consistent train and test sets to evaluate all methods, we analyzed the integrity of the databases against each other with relation to the occurrence of duplicated sequences.

**Figure 1.**
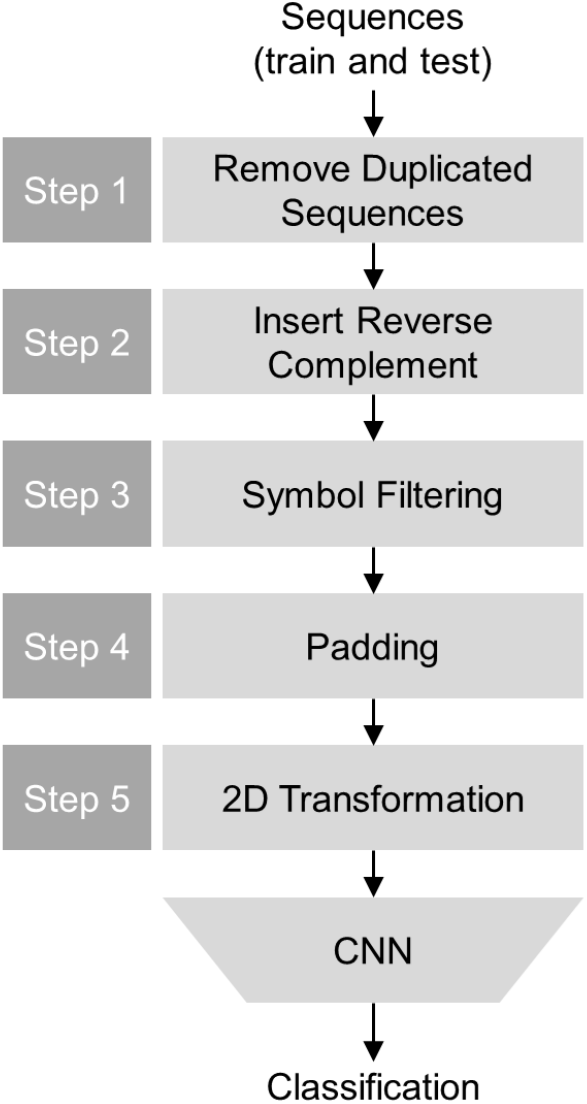
Pipeline of TERL. The input consists of train and test sets of sequences. Step 1 removes duplicated sequences from the databases to build the sets. Step 2 inserts in the sets the reverse-complemented counterparts of the sequences. Step 3 changes all symbols to upper case and map any different symbol than {*A,C,G,T,N*} to *N*. Step 4 pads the sequences with symbol *B* to adjust them to same length as the longest sequence. Step 5 applies the transformation from one-dimensional nucleic acid sequences to two-dimensional space data. On the CNN step the twodimensional space data are applied to the CNN to train the model and test it.

We analyzed seven databases and found duplicated sequences that are classified in both databases as the same class and also as different classes. In this preprocessing step, we examine if every sequence presents copies in other databases. For every copy found we analyze if the sequences represent the same class. If they represent the same class, we keep the sequence from the database with the least number of samples for the respective class and remove the other sequence. If the sequences do not represent the same class, both sequences are removed since there is a divergence between databases with relation to the classes of the sequences. We remove these sequences to avoid overfitting and misleading performance results since the samples could represent different classes in train and test sets. In our pipeline, this preprocessing step is referred as step 1 in Figure 1.

Since DNA sequences have their reverse-complemented counterparts, we defined a step to insert the reverse-complemented counterpart of each training sequence in the training data set. This adds information to the network to train it to recognise possible reverse-complemented counterparts. This step is referred as step 2 in Figure 1.

TE sequences are usually codified with the symbols {*A, C, G,T,N*} (four nucleotides and one unidentified nucleotide symbol *N*). TE sequences may also present different symbols than {*A, C, G, T, N*} to represent uncertainties during sequencing. These uncertain symbols are preprocessed and altered to *N* to simplify the representation of the sequences. In our method, this step is referred as step 3 in Figure 1.

All sequences are padded to the length of the longest sequence in the data sets. The sequences can also be padded to a longer length value that can allow the network to accept sequences longer than the ones from the data sets. In this preprocessing step, background symbols (*B*) are inserted in the sequences to normalize the length of all sequences. This new symbol is used since the padding of a sequence is a new information and does not correspond to any nucleotide or unidentified nucleotide. In our pipeline, this preprocessing step is referred as step 4 in Figure 1.

To apply these sequences into the CNN, the input data must be numerical and normalized. Our approach uses one-hot encoding vectors to transform TE sequences into a matrix of ones and zeros. This transformation preserves information contained in the raw sequences regarding the positions of the nucleotides. Each nucleotide is mapped to value 1 in its specific row and the other rows are filled with value 0. The columns represent the nucleotide sequence of the transposons. This preprocessing step is referred as step 5 in Figure 1.

Each line of the one-hot encoding matrix represents one specific symbol from the set {*A, C, G, T, N, B*} (being *A, C, G* and *T* the four nucleotides, N the unknown nucleotide and B the padding symbol). An example of the one-hot encoded vector for a TE sequence is shown in Figure 2.

**Figure 2.**
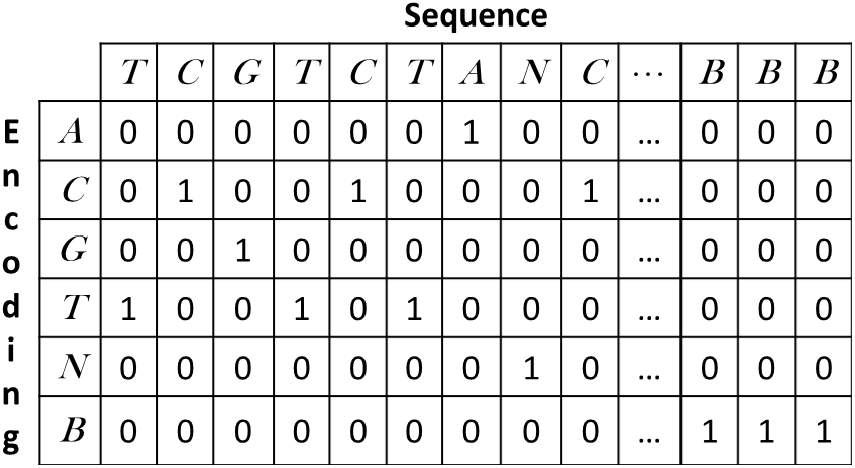
Two-dimensional space transformation by one-hot encoding example. The columns represent the symbols of the sequence, which can be any element from the set {A, *C, G, T, N, B*}. Each line represents a specific symbol and for each column, the line representing the same symbol as the column has value 1 and for that column all other lines have value 0.

After all these preprocessing steps, the deep CNN model can be trained with the sequences from the train set and its performance can be calculated with the sequences from the test set.

This approach is generic and can be applied to classify any biological sequence. The CNN architecture is arbitrary and can be designed to improve performance for the type of sequence that will be classified.

In our approach, the height of the convolution filters of the first convolution layer is equal to the number of encoding symbols (i.e. the height of the first convolution layers is equal to the number of rows of the one-hot encoding matrix shown in Figure 2), thus the convolution filter comprises all the information of a delimited region.

This height is defined since the two-dimensional space transformation represents each nucleotide of the sequence with values for all symbols from the set {*A, C, G,T, N, B*}. The following pooling and convolution layers have one dimension due to the height reduction of the first convolution layer.

### Experiments

We have designed six experiments to compare the performance of TERL against other methods and to verify its performance on classifying TE sequences into superfamily level. These experiments consist in classifying sequences from five designed data sets. We also verify how well TERL can generalize compared to other methods.

The proposed pipeline was implemented on Python programming language. All CNN models were implemented with Tensorflow version 1.13. The tests were performed on a computer with processor Intel(R) Xeon(R) E5-2620 v3 @ 2.4 GHz, DDR4 SDRAM memory of 24 GB @ 2133 MHz and NVIDIA TITAN V GPU.

#### Data Sets Descriptions

The data sets were built using sequences from seven TEs databases. The databases used are: RepBase [15], DPTEdb [19], SPTEdb [27], PGSB PlantsDB [24], RiTE database [6], TREP [25] and TEfam.

RepBase is one of the most used TE database and it is composed by sequences from more than 100 organisms, including sequences from animals, plants and fungi genomes.

DPTEdb, PGSB PlantsDB, RiTE database and SPTEdb are plant-specific TE databases composed by sequences from eight, 18, 12 and three genomes respectively.

TEfam is a TE database composed by two mosquitoes species genomes, *Aedes aegypti* and *Anopheles gambiae.* TREP is a database composed by sequences from 49 organisms, including plants, animals, fungi and bacteria.

We have defined five data sets to evaluate the performance of TERL and conduct the comparisons with other methods. Data set 1 consists of the Copia, Gypsy, Bel-Pao, ERV, L1, Mariner and hAT superfamilies of RepBase sequences and non-TE sequences that were generated by applying shuffling in the sequences from these superfamilies (see Table 2). The amount of training and testing sequences was defined to 80% (1364) and 20% (341) respectively of the class with least total sequences (i.e. L1). The other classes were undersampled to that amount of 1364 training and 341 testing samples.

**Table 2.** Sequences distribution of data set 1. Train and test sets are composed by RepBase sequences and non-TE sequences. Sets are undersampled to 1364 training sequences and 341 testing ones.

| Superfamily | Sequences | Train | Test |
| --- | --- | --- | --- |
| Copia | 7025 | 1364 | 341 |
| Gypsy | 11170 | 1364 | 341 |
| Bel-Pao | 1871 | 1364 | 341 |
| ERV | 4439 | 1364 | 341 |
| L1 | 1705 | 1364 | 341 |
| hAT | 3044 | 1364 | 341 |
| Tc1-Mariner | 2589 | 1364 | 341 |
| Non-TE | - | 1364 | 341 |

All non-TE sequences used in this paper were generated by sampling the sequences from other classes and applying shuffling to them.

Data set 2 consists of the orders (LTR and LINE), class (Class II) and non-TE sequences (see Table 3). The amount of training and testing samples were defined to 80% (2280) and 20% (570) of the class with least total sequences (i.e. LINE). The other classes were undersampled to the same amount of 2280 training sequences and 570 testing ones.

**Table 3.** Sequences distribution of data set 2. Train and test sets are composed by RepBase sequences and non-TE sequences. Sets are undersampled to 2280 training sequences and 570 testing ones.

| Order | Total | Train | Test |
| --- | --- | --- | --- |
| LTR | 24505 | 2280 | 570 |
| LINE | 2850 | 2280 | 570 |
| Class II | 9623 | 2280 | 570 |
| Non-TE | - | 2280 | 570 |

Data set 3 is composed by the Copia, Gypsy, Bel-Pao, ERV, L1, SINE, Tc1-Mariner, hAT, Mutator, PIF-Harbinger and CACTA superfamilies sequences from the seven databases and non-TE sequences (see Table 4). Each class was undersampled to 80% (2177) and 20% (545) of the total sequences of the class with least total sequences (i.e. Bel-Pao). The amount of sequences for each class in each database is shown in Table 4.

**Table 4.** Sequences distribution of data set 3. Train and test sets are composed by sequences from all seven databases and non-TE sequences. Sets are undersampled to 2177 training sequences and 545 testing ones.

| Superfamily | DPTE | PGSB | REPBAS | RiTE | SPTE | TEfam | TREP | Total | Train | Test |
| --- | --- | --- | --- | --- | --- | --- | --- | --- | --- | --- |
| Copia | 3820 | 3972 | 7025 | 13596 | 2764 | 323 | 418 | 31918 | 2177 | 545 |
| Gypsy | 5756 | 7220 | 11170 | 63457 | 6562 | 454 | 525 | 95144 | 2177 | 545 |
| Bel-Pao | 211 | 0 | 1871 | 2 | 144 | 494 | 0 | 2722 | 2177 | 545 |
| ERV | 583 | 0 | 4439 | 325 | 104 | 0 | 0 | 5451 | 2177 | 545 |
| L1 | 1299 | 0 | 1705 | 737 | 278 | 114 | 3 | 4136 | 2177 | 545 |
| SINE | 0 | 191 | 239 | 3072 | 0 | 0 | 0 | 3502 | 2177 | 545 |
| Tc1-Mariner | 50 | 322 | 2589 | 19898 | 0 | 32 | 375 | 23266 | 2177 | 545 |
| hAT | 96 | 76 | 3044 | 3700 | 40 | 42 | 25 | 7023 | 2177 | 545 |
| Mutator | 23 | 212 | 1380 | 99498 | 8 | 4 | 163 | 101288 | 2177 | 545 |
| PIF-Harbinger | 91 | 86 | 1128 | 17878 | 11 | 12 | 109 | 19315 | 2177 | 545 |
| CACTA | 0 | 454 | 718 | 9163 | 0 | 0 | 324 | 10659 | 2177 | 545 |
| Non-TE | - | - | - | - | - | - | - | - | 2177 | 545 |

Data set 4 consists of orders (LTR, LINE and SINE) and class (Class II) from the seven databases and non-TE sequences (see Table 5). Each class was undersampled to 80% (3158) and 20% (790) of the total sequences of the class with least total sequences (i.e. SINE). The amount of sequences for each class in each database is also shown in Table 5.

**Table 5.** Sequences distribution of data set 4. Train and test sets are composed by sequences from all seven databases and non-TE sequences. Train set is undersampled to 3158 sequences per class and test set to 790.

| Order | DPTE | PGSB | REPBAS | RiTE | SPTE | TEfam | TREP | Total | Train | Test |
| --- | --- | --- | --- | --- | --- | --- | --- | --- | --- | --- |
| LTR | 10370 | 11192 | 24505 | 77380 | 9574 | 1271 | 943 | 135235 | 3158 | 790 |
| LINE | 1299 | 470 | 2850 | 784 | 278 | 368 | 8 | 6057 | 3158 | 790 |
| SINE | 0 | 191 | 685 | 3072 | 0 | 0 | 0 | 3948 | 3158 | 790 |
| Class II | 260 | 1150 | 9623 | 150142 | 59 | 128 | 996 | 162358 | 3158 | 790 |
| Non-TE | - | - | - | - | - | - | - | - | 3158 | 790 |

Training sequences of the data set 5 were sampled from orders sequences from RepBase database and undersampled to 2850, which is the total sequence of the class with least total sequences on RepBase (i.e. LINE). The testing sequences of the data set 5 were sampled from orders sequences from the six remaining databases and undersampled to 570, which is 20% of the training samples for each class. This data set is composed by sequences from orders (LTR and LINE), class (Class II) and non-TE sequences (see Table 6).

**Table 6.**
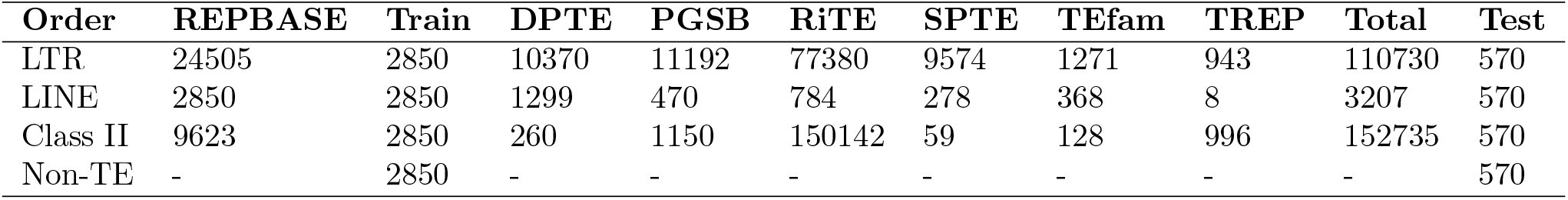
Sequences distribution of data set 5. Train set is composed by sequences from RepBase and non-TE sequences. Test set is composed by sequences from the six remaining databases and non-TE sequences. Train set is undersampled to 2850 sequences per class and test set to 570.

The classes are balanced to avoid over-representation of some classes, which can negatively affect model training [17] [2]. The under-sampling consists of grouping sequences by class and randomly selecting the amount of sequences of the least represented class from all classes. The sequences are selected based on an uniform distribution, thus the sequences from databases that present more sequences for a specific class have more probability to be selected. This is necessary due to lack of sequences for some specific classes in some databases.

#### Scenarios

We propose six experiments to verify the performance of TERL on the classification of: TE sequences into superfamily, order and class levels. All experiments use the CNN: architecture shown in Figure 3. The architecture is composed of 9 layers which are: described in Table 7.

**Figure 3.**
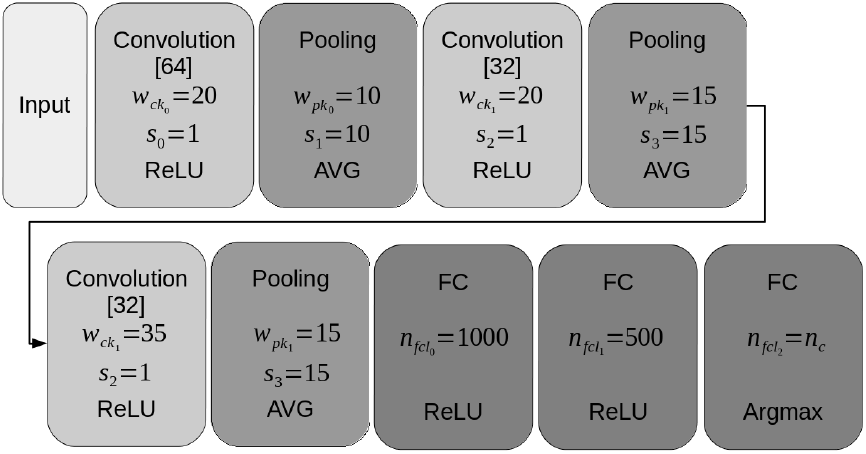
CNN architecture. Each block represents a layer. The type of the layer is described on the top of each block. The numbers inside brackets are the number of feature maps. *w* describes the width, *s* the strides and *n* the neurons of the layers. The activation function is described on the bottom of each block.

**Table 7.** Description of each layer of the CNN architecture used on TERL. Column Filters/Neurons describes the number of filters for convolution layers and neurons for fully connected layers.

| Layer | Width | Height | Stride | Filters/Neurons | Function |
| --- | --- | --- | --- | --- | --- |
| Convolution | 20 | 6 | 1 | 64 | ReLU |
| Pooling | 10 | 1 | 10 | - | Average |
| Convolution | 20 | 1 | 1 | 32 | ReLU |
| Pooling | 15 | 1 | 15 | - | Average |
| Convolution | 35 | 1 | 1 | 32 | ReLU |
| Pooling | 15 | 1 | 15 | - | Average |
| Fully connected | - | - | - | 1000 | ReLU |
| Fully connected | - | - | - | 500 | ReLU |
| Fully connected | - | - | - | $N_c$ | Softmax |

The loss function of the trained models is softmax cross entropy. The learning algorithm used was Adam with learning rate 0.001 and l2 regulation of 0.001. Each experiment is executed for 100 epochs with train and test batch of 32 samples.

Experiments 1 and 2 analyze the performance of TERL on classifying RepBase sequences into superfamily and order levels respectively. The data sets 1 and 2 are used in experiments 1 and 2, respectively. On experiment 2, the results of the classification into order level are compared to the results of TEclass and PASTEC, since none of these methods classifies the sequences into superfamily level.

In the experiment 2 we verify the ability of each method to classify curated sequences from the reference database for eukaryotic TEs sequences, RepBase. The sequences from RepBase usually do not present long unidentified inter-contig regions. These regions have no information to present to the network and can be considered as noise during the training phase.

Experiments 3 and 4 analyze the performance of TERL on classifying sequences of the data sets 3 and 4, respectively. Experiment 3 classifies the sequences into superfamily level and experiment 4 into order level. The purpose of these experiments is to analyze the ability of TERL to classify sequences from different databases (i.e. different organisms and different sequencing qualities).

Different databases can present sequences with different sequencing qualities, not well-annotated sequences and unidentified inter-contig regions, which could be considered as noise in the classification process. When sequences are not well-annotated, the performance of the classification algorithms can drop since there can be sequences with wrong label in the training and test sets. The results obtained on experiment 4 are compared with those obtained by TEclass and PASTEC.

Experiment 5 analyzes the performance of TERL when using only RepBase sequences to train the model and testing it with sequences from the six remaining databases (i.e. data set 5). The sequences are classified into order level and the goal of this experiment is to analyze the generalization of the methods since RepBase sequences are curated, well-annotated and present few unknown regions (i.e., regions with symbols different than {*A,C,G,T*}, and the sequences from other databases can present unidentified regions and may not be well-annotated. The results are compared with those obtained by TEclass and PASTEC.

Every experiment consists of retraining the CNN used on TERL to learn the patterns of the classes used in the respective experiment.

Since PASTEC method is a pseudo agent system and can be used with different auxiliary files, on Experiment 6 we analyze the performance of PASTEC in case each homology search is turned off and compare these results with those obtained by TERL. In this experiment we test the classification of data sets 2, 4 and 5.

This experiment also verifies the ability of PASTEC to classify the sequences without any prior information, as TERL does not need any prior information, just the sequences.

## Results and Discussion

We evaluate TERL on the classification of five data sets (2 superfamily and 3 order levels data sets) and compare the results with PASTEC and TEclass on the classification of test sequences from the 3 order level data sets. Initially, we evaluate the performance of TERL in the first experiment, using the data set 1 and classifying the sequences considering seven superfamilies from RepBase and non-TE sequences.

According to the results presented in Figure 4 and on Table 8, it is clear that using CNNs to classify TE sequences, TERL is capable of learning how to recognize the patterns of these sequences and classify them into superfamilies since it obtained a macro mean accuracy of 96.4% and F1-score of 85.8%. Few methods on the literature present results for TE classification into the superfamily level and with superfamilies from different orders. The metrics for multi-class classification are calculated based on the equations on [23].

**Figure 4.**
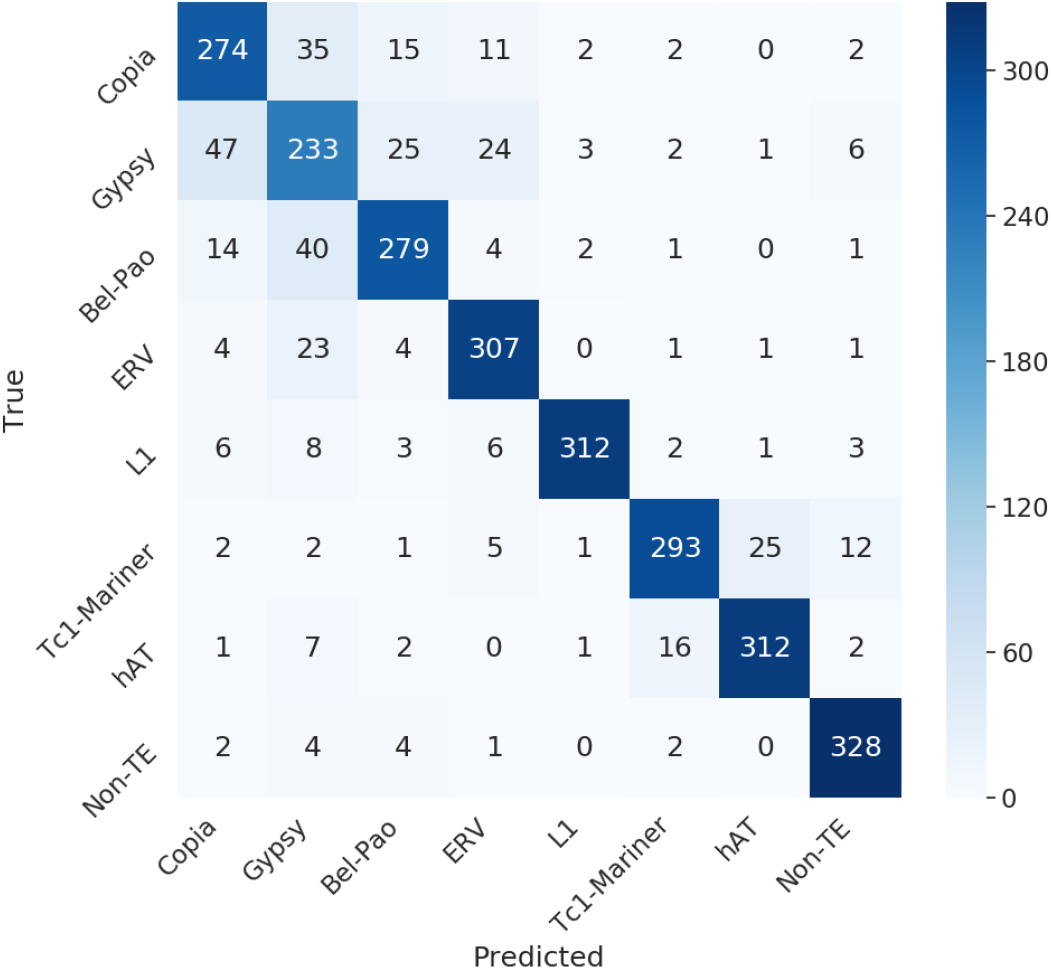
Confusion matrix obtained on experiment 1 on the classification of data set 1 into superfamily level by TERL using a CNN with the architecture described on Table 7.

**Table 8.**
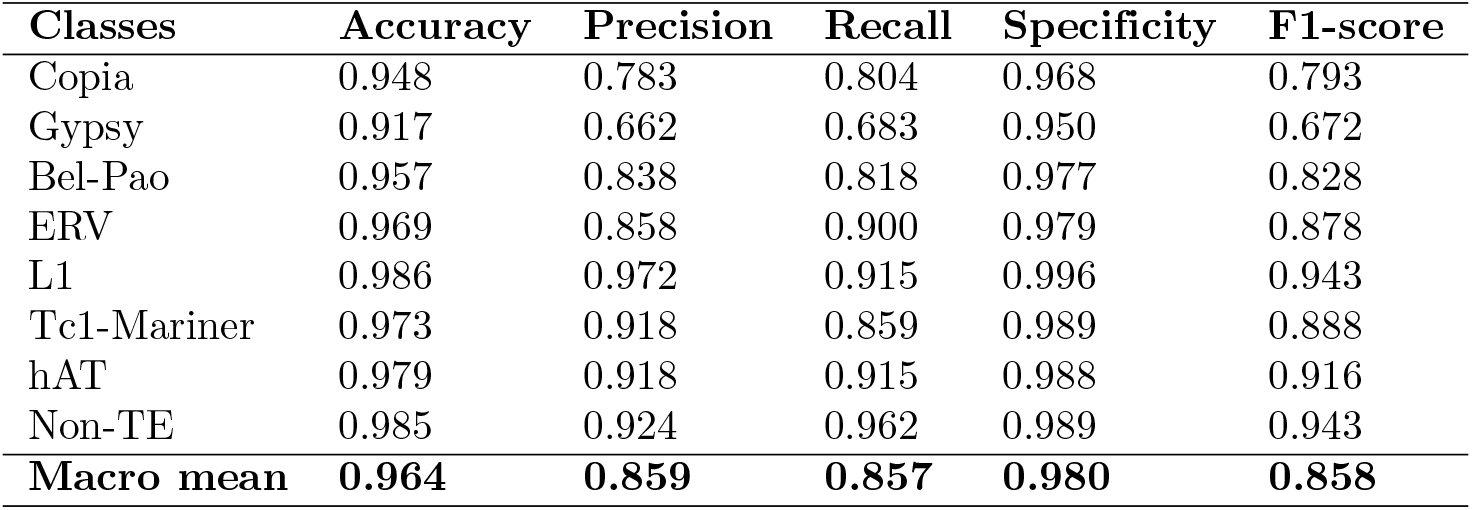
Metrics obtained on experiment 1 on the classification of sequences from data set 1 by TERL using the CNN with the arhitecture described on Table 7.

| Classes | Accuracy | Precision | Recall | Specificity | F1-score |
| --- | --- | --- | --- | --- | --- |
| Copia | 0.948 | 0.783 | 0.804 | 0.968 | 0.793 |
| Gypsy | 0.917 | 0.662 | 0.683 | 0.950 | 0.672 |
| Bel-Pao | 0.957 | 0.838 | 0.818 | 0.977 | 0.828 |
| ERV | 0.969 | 0.858 | 0.900 | 0.979 | 0.878 |
| L1 | 0.986 | 0.972 | 0.915 | 0.996 | 0.943 |
| Tc1-Mariner | 0.973 | 0.918 | 0.859 | 0.989 | 0.888 |
| hAT | 0.979 | 0.918 | 0.915 | 0.988 | 0.916 |
| Non-TE | 0.985 | 0.924 | 0.962 | 0.989 | 0.943 |
| **Macro mean** | **0.964** | **0.859** | **0.857** | **0.980** | **0.858** |

Figure 4 presents a confusion matrix indicating that for each superfamily the false negatives are more concentrated in the superfamilies from the same order. This means that the CNN was able to learn how to recognize the orders’ patterns as well.

Analyzing the results (Figure 4 and Table 8), the Gypsy superfamily was the class that presented lower results and more confusion than the others. The method was not able to learn how to correctly recognize the sequences from the Gyspsy superfamily, given the classification distribution on the Gypsy row and column indicates that some real Gypsy sequences were wrongly classified as other LTR superfamilies (Bel-Pao, Copia, ERV) and Bel-Pao, Copia and ERV superfamilies sequences were classified as Gypsy.

Tc1-Mariner and hAT superfamilies, which are superfamilies from the same order (TIR), also presented some higher confusion between them. This indicates that the method did not learn how to properly recognize them, but learned that these two superfamilies sequences are related and close to each other.

We also compare the results of TERL with those of other literature methods, PASTEC and TEclass, in the experiment 2, using the data set 2 (see Figure 5 and Table 9. Based on the results, TERL outperforms TEclass considering all metrics (accuracy, precision, recall, specificity and F1-score). In this experiment TERL could not surpass the results of PASTEC, but obtained similar results, as the difference between the results were 0.023, 0.048, 0.047, 0.015 and 0.047 for accuracy, precision, recall, specificity and F1-score respectively.

**Figure 5.**
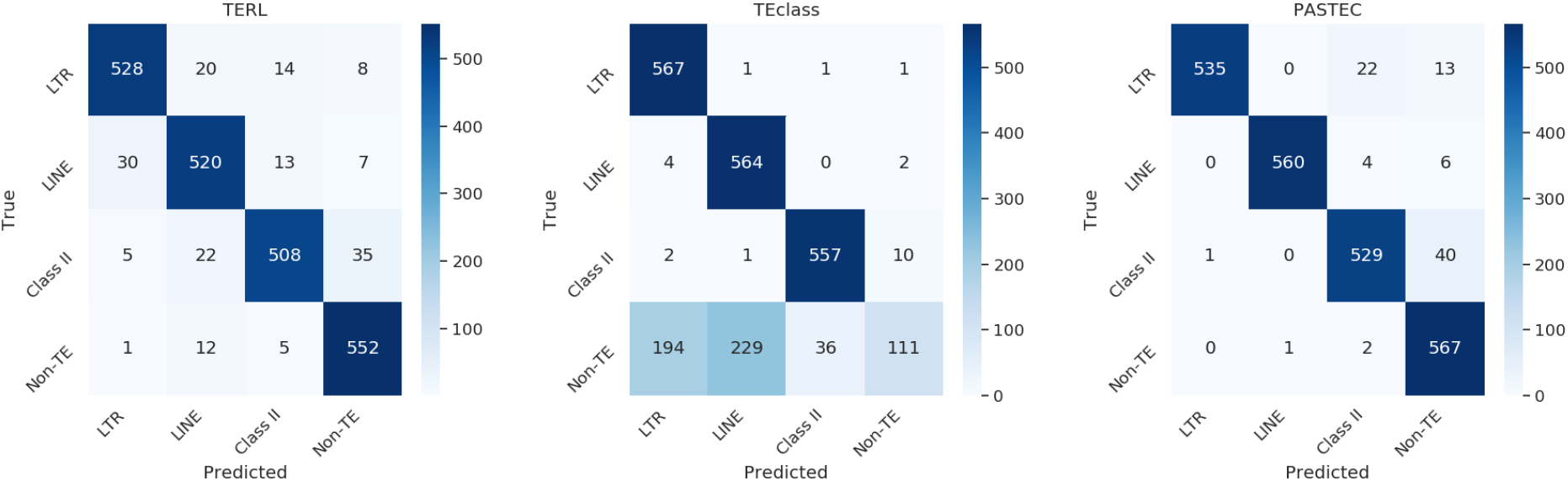
Confusion matrix obtained on experiment 2 on the classification of data set 2 into order level by TERL, TEclass and PASTEC.

**Table 9.** Metrics obtained on experiment 2 on the classification of sequences from data set 2 by TERL using the CNN with the arhitecture described on Table 7, PASTEC and TEclass.

| Class | Method | Accuracy | Precision | Recall | Specificity | F1-score | Test time |
| --- | --- | --- | --- | --- | --- | --- | --- |
| LTR | TERL | 0.947 | 0.895 | 0.896 | 0.965 | 0.895 |  |
|  | PASTEC | **0.984** | **0.998** | 0.939 | **0.999** | **0.967** |  |
|  | TEclass | 0.911 | 0.739 | **0.995** | 0.883 | 0.848 |  |
| LINE | TERL | 0.961 | 0.910 | 0.937 | 0.969 | 0.923 |  |
|  | PASTEC | **0.995** | **0.998** | 0.982 | **0.999** | **0.990** |  |
|  | TEclass | 0.896 | 0.709 | **0.989** | 0.865 | 0.826 |  |
| Class II | TERL | 0.952 | 0.935 | 0.868 | 0.9797 | 0.900 |  |
|  | PASTEC | 0.970 | **0.950** | 0.928 | **0.984** | 0.939 |  |
|  | TEclass | **0.978** | 0.938 | **0.977** | 0.978 | **0.957** |  |
| Non-TE | TERL | 0.969 | **0.921** | 0.957 | 0.973 | 0.938 |  |
|  | PASTEC | **0.973** | 0.906 | **0.995** | 0.965 | **0.948** |  |
|  | TEclass | 0.793 | 0.895 | 0.195 | **0.992** | 0.320 |  |
| Macro mean | TERL | 0.957 | 0.915 | 0.914 | 0.972 | 0.915 | **1.044s** |
|  | PASTEC | **0.980** | **0.963** | **0.961** | **0.987** | **0.962** | 433m 11.342s |
|  | TEclass | 0.895 | 0.820 | 0.789 | 0.930 | 0.804 | 2m 54.171s |

Both PASTEC and TEclass use homology search in RepBase sequences database. PASTEC searches in nucleotides, amino acids and HMMs profiles databases to produce its outputs, thus there is a possibility of overfitting in the results of this method, since the complete exclusion of test set sequences in the auxiliary files could not be verified and guaranteed.

The computation time was also evaluated in this experiment, and TERL can classify the same amount of sequences much faster, by using GPU. All experiments were executed on a computer with CPU Intel Xeon E5-2620 v3 2.4 GHz x 4, 26 GB of DDR4 RAM 2133 MHz, GPU NVIDIA TITAN V with 12 GB GDDR5 graphic memory and 5120 CUDA cores.

As Table 9 shows, TERL is by far the fastest method evaluated. Time values for PASTEC is related to the time of the longest batch because we threaded its execution with batchs of the test set. The batches were divided by class, which means the longest class took more than 433 minutes. TEclass is also faster than PASTEC, but not as fast as TERL.

Experiment 3 analyzes the performance of TERL on the classification of sequences from all seven databases into 11 superfamilies. Each database presents sequences from several species and can present distinct sequencing qualities, thus this experiment tests the ability of the methods to be generalized.

Sequences from some databases can present long unidentified inter-contig regions, which consist of long sequences of symbols N and may present a challenge to pattern recognition methods since they need to learn how to recognize these patterns with these unidentified regions as noise.

Based on the results shown in Figure 6 and on Table 10, TERL was capable to correctly classify sequences into 11 superfamilies of the data set 3. We obtained a macro mean accuracy, precision, recall, specificity and F1-score of 0.95, 0.711, 0.701, 0.973 and 0.706 respectively. As can also be observed (Table 10 and Figure 6), the multi-class accuracy metric can be misleading, given we obtained an accuracy of 0.95 compared with the precision, recall and F1-score of 0.711, 0.701 and 0.706 respectively.

**Figure 6.**
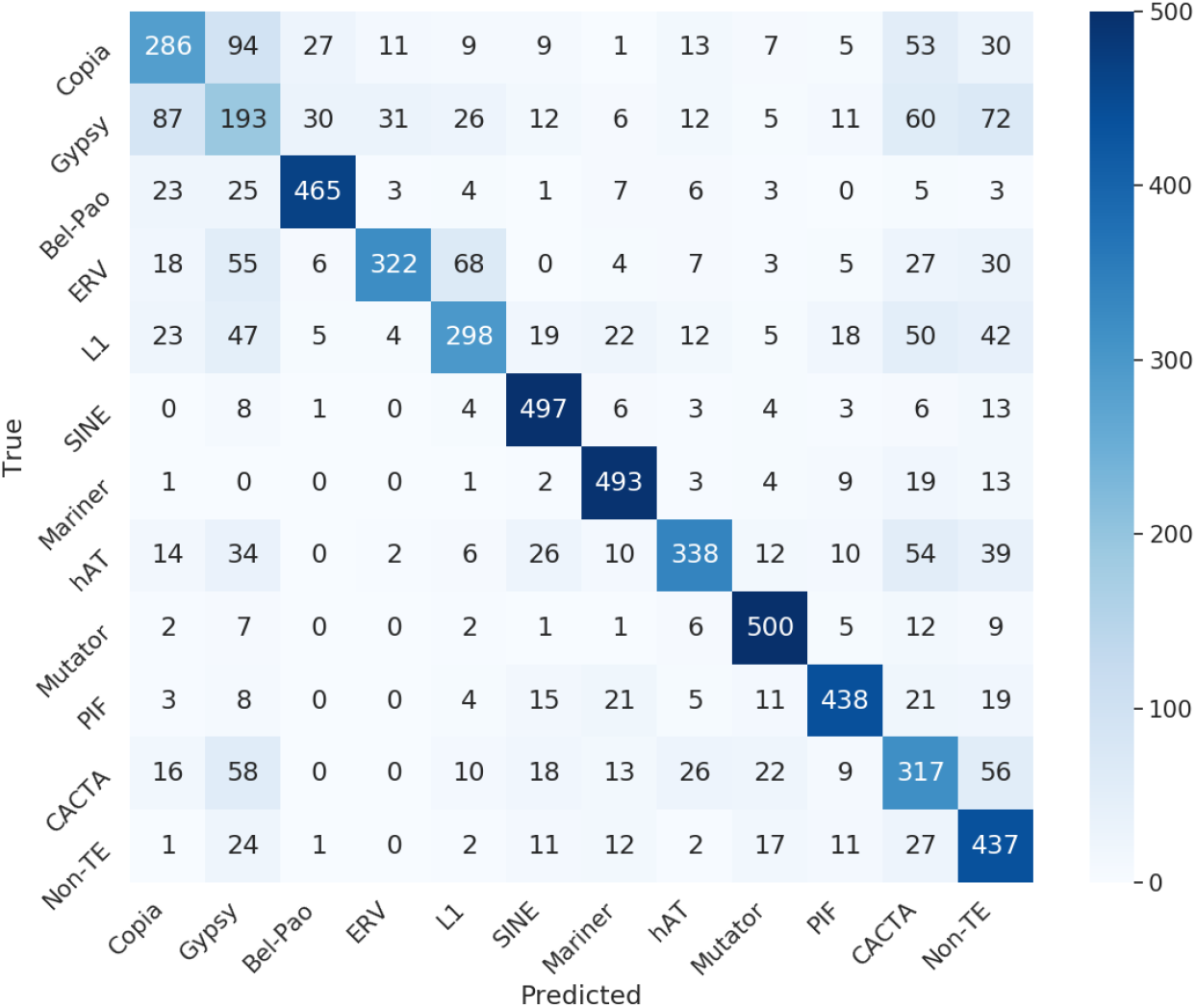
Confusion matrix obtained on experiment 3 on the classification of data set 3 into superfamily level by TERL using a CNN with the architecture described on Table 7.

**Table 10.** Metrics obtained on experiment 3 on the classification of sequences from data set 3 by TERL using the CNN with the arhitecture described on Table 7.

| Class | Accuracy | Precision | Recall | Specificity | F1-score |
| --- | --- | --- | --- | --- | --- |
| Mutator | 0.979 | 0.843 | 0.917 | 0.984 | 0.879 |
| Mariner | 0.976 | 0.827 | 0.905 | 0.983 | 0.864 |
| Copia | 0.932 | 0.603 | 0.525 | 0.969 | 0.561 |
| CACTA | 0.914 | 0.487 | 0.582 | 0.944 | 0.530 |
| ERV | 0.958 | 0.863 | 0.591 | 0.991 | 0.702 |
| L1 | 0.941 | 0.687 | 0.547 | 0.977 | 0.609 |
| Gypsy | 0.891 | 0.349 | 0.354 | 0.940 | 0.352 |
| PIF | 0.970 | 0.836 | 0.804 | 0.986 | 0.819 |
| hAT | 0.954 | 0.781 | 0.620 | 0.984 | 0.691 |
| Random | 0.934 | 0.573 | 0.802 | 0.946 | 0.668 |
| SINE | 0.975 | 0.813 | 0.912 | 0.981 | 0.860 |
| Bel-Pao | 0.977 | 0.869 | 0.853 | 0.988 | 0.861 |
| **Macro mean** | **0.950** | **0.711** | **0.701** | **0.973** | **0.706** |

As the results obtained on experiment 1, the confusion matrix shows that our approach was able to recognize the patterns that define the sequences from the LTR order since the major error zone in the graph is in the region of the LTR superfamilies. These sequences tend to be very similar to each other, as presenting almost the same structure and sometimes the same protein domains. The method also wrongly predicted sequences as non-TE sequences.

The Gypsy superfamily was again the superfamily with lower metrics, showing a confusion and difficulty to learn how to correctly recognize the patterns of these sequences.

Compared to the Figure 4 and Table 8, this classification does not present the same performance, which can be due to the insertion of sequences from different organisms, bad annotated sequences and sequences with long unidentified regions.

To compare the results of the classification with those obtained by TEclass and PASTEC, considering the data set 4, the network is retrained to classify the sequences into order level. Figure 7 and Table 11 present the confusion matrix and the metrics obtained by the methods on this classification.

**Figure 7.**
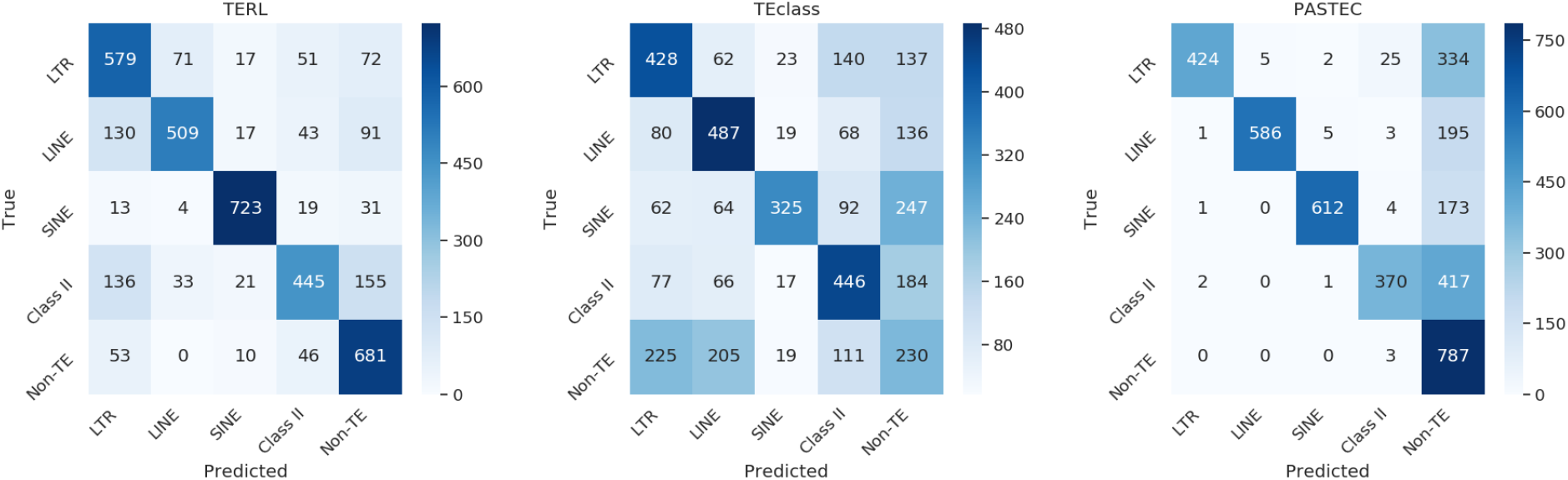
Confusion matrix obtained on experiment 4 on the classification of data set 4 into order level by TERL using a CNN with the architecture described on Table 7, TEclass and PASTEC.

According to the obtained results, TERL is once again more efficient than TEclass to classify sequences from all databases, which is a more realistic scenario for TE classification. TERL outperformed TEclass in all metrics on Table 11. TERL obtained 0.893, 0.745, 0.732, 0.933 and 0.739 of accuracy, precision, recall, specificity and F1-score respectively.

**Table 11.**
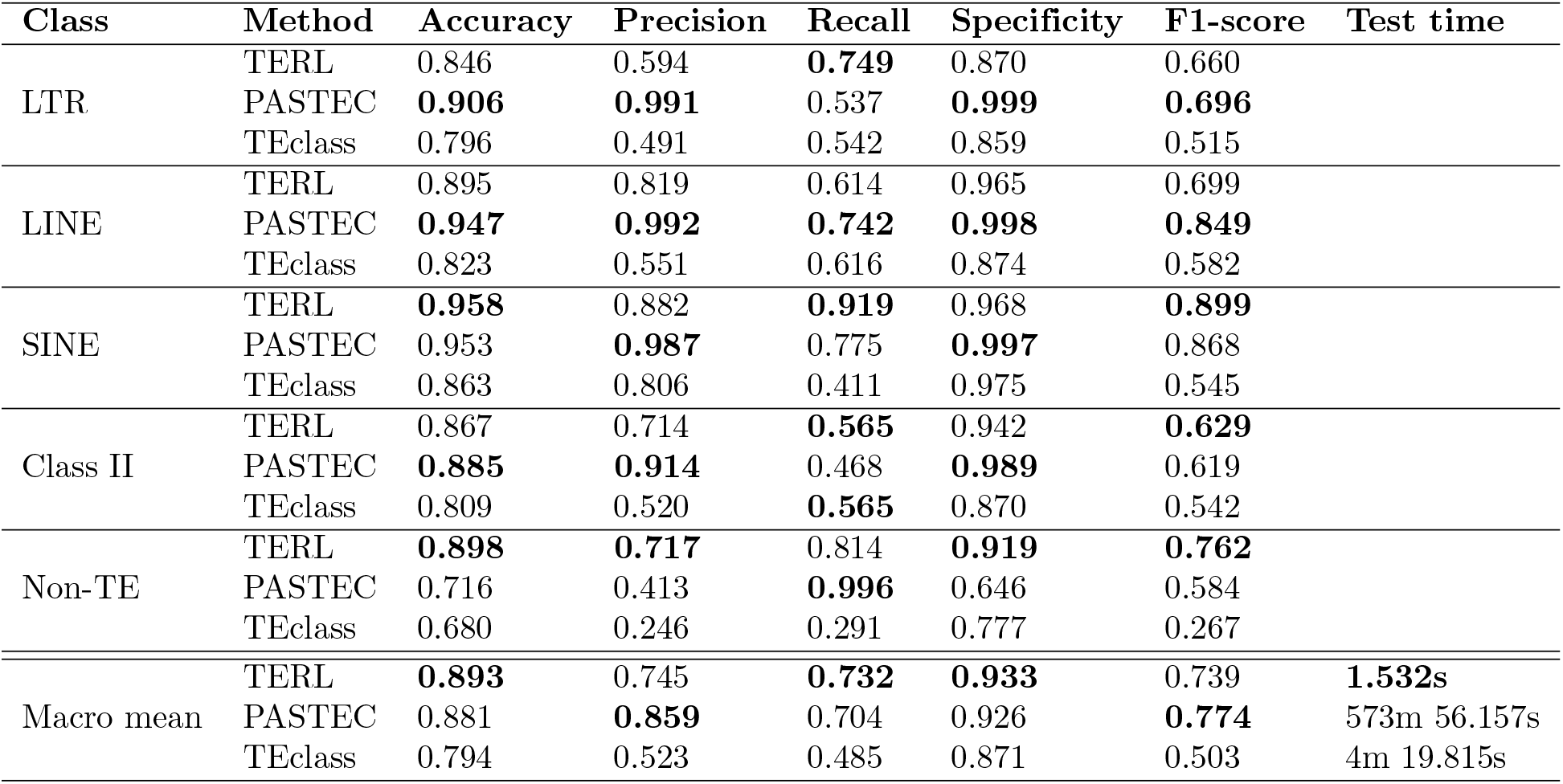
Metrics obtained on experiment 4 on the classification of sequences from data set 4 by TERL using the CNN with the arhitecture described on Table 7, PASTEC and TEclass.

In this experiment TERL was capable to surpass the accuracy, recall and specificity metrics obtained by the PASTEC method. The other metrics were similar to those obtained by PASTEC since the difference between precision and F1-score was 0.014 and 0.035 respectively.

Figure 7 shows that TERL obtained better results, given the best true positive obtained by TEclass was only 487 sequences against the best true positive of TERL which was 723 sequences.

The classification obtained by TEclass was also very sparse throughout the classes and this method was clearly not capable to predict non-TE sequences.

Although PASTEC obtained some metrics close to the ones obtained by TERL, the method predicted a great number of sequences as non-TE sequences for all classes, leading to an accuracy and precision of non-TE sequences of 0.716 and 0.413 respectively. PASTEC only classifies the sequences when the evidence is above a threshold; otherwise it classifies them as non-TE sequences.

In this experiment PASTEC method could be presenting overfitting again by using RepBase sequences as auxiliary file for the homology searches and the HMMs profile file can present some of the protein domains of the sequences in the test set.

Table 11 also shows that TERL classified the sequences in about 1.53 seconds, against 3.44 × 10^4^ and 2.59 × 10^2^ seconds for PASTEC and TEclass, respectively. Again TERL is the fastest method evaluated. TEclass again is faster than PASTEC, but not as fast as TERL. In this experiment PASTEC took more than 573 minutes to finish the execution of one of the batchs. This time is the longest time took by a class batch.

To provide a fare comparison, the experiment 5 analyzes the capability of the methods to classify TEs into order level by training with RepBase sequences and testing with sequences from all other databases (i.e. data set 5). Figure 8 shows the confusion matrices obtained by TERL, TEclass and PASTEC. Table 12 presents the metrics obtained by the methods.

**Figure 8.**
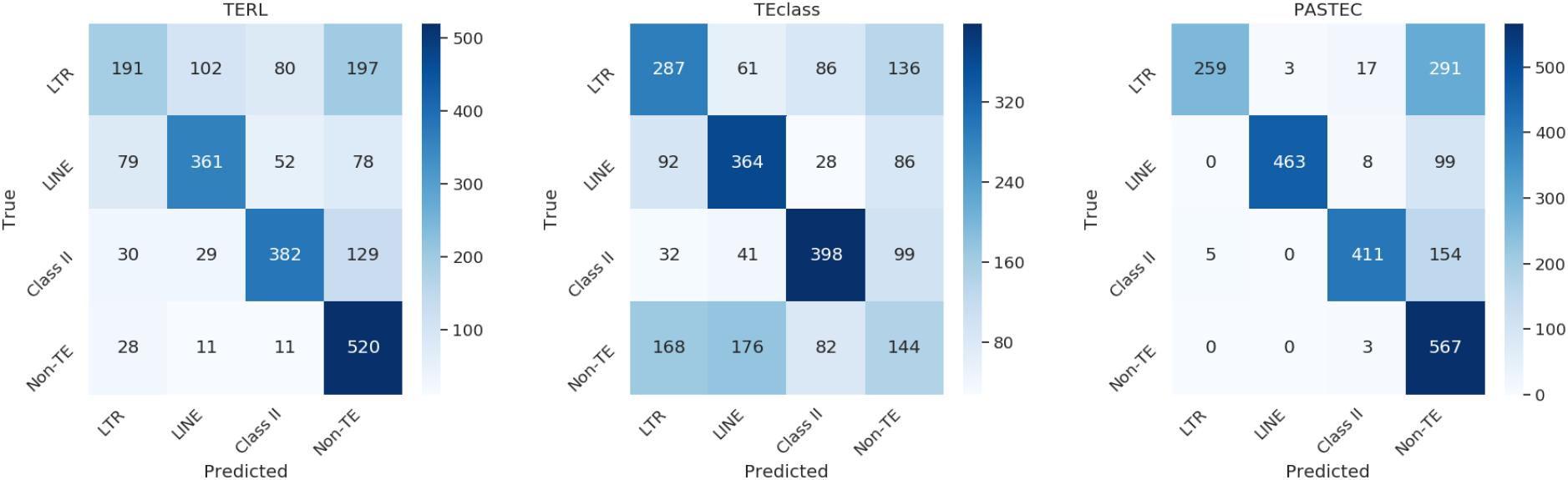
Confusion matrix obtained on experiment 5 on the classification of data set 5 into order level by TERL using the CNN with architecture described on Table 7, TEclass and PASTEC.

**Table 12.** Metrics obtained on experiment 5 on the classification of sequences from data set 5 by TERL using the CNN with the arhitecture described on Table 7, PASTEC and TEclass.

| Class | Method | Accuracy | Precision | Recall | Specificity | F1-score | Test time |
| --- | --- | --- | --- | --- | --- | --- | --- |
| LTR | TERL | 0.768 | 0.564 | 0.363 | 0.903 | 0.436 |  |
|  | PASTEC | **0.861** | **0.981** | 0.454 | **0.997** | **0.621** |  |
|  | TEclass | 0.748 | 0.496 | **0.504** | 0.829 | 0.500 |  |
| LINE | TERL | 0.820 | 0.669 | 0.570 | 0.904 | 0.613 |  |
|  | PASTEC | **0.952** | **0.994** | **0.812** | **0.998** | **0.894** |  |
|  | TEclass | 0.788 | 0.567 | 0.639 | 0.837 | 0.601 |  |
| Class II | TERL | 0.826 | 0.649 | 0.683 | 0.874 | 0.663 |  |
|  | PASTEC | **0.918** | **0.936** | **0.721** | **0.984** | **0.815** |  |
|  | TEclass | 0.839 | 0.670 | 0.698 | 0.885 | 0.684 |  |
| Non-TE | TERL | **0.829** | **0.613** | 0.870 | **0.815** | **0.718** |  |
|  | PASTEC | 0.760 | 0.510 | **0.995** | 0.682 | 0.675 |  |
|  | TEclass | 0.672 | 0.310 | 0.253 | 0.812 | 0.278 |  |
| Macro mean | TERL | 0.811 | 0.624 | 0.621 | 0.874 | 0.623 | **1.097s** |
|  | PASTEC | **0.873** | **0.855** | **0.746** | **0.915** | **0.797** | 275m 7.478s |
|  | TEclass | 0.762 | 0.511 | 0.523 | 0.841 | 0.517 | 2m 45.858s |

The results in Figure 8 and Table 12 show that TERL surpassed TEclass. TEclass is not able to recognize non-TE sequences since the non-TE row presents almost evenly distributed classifications throughout the classes.

PASTEC method obtained the best results in this experiment, except for test time, which TERL was about 1.50 × 10^4^ times faster. The confusion among other orders is very low and the method still tends to predict non-TE sequences when the evidence is not sufficient. The difference between the results obtained by PASTEC and the ones obtained by TERL was 0.062, 0.231, 0.125, 0.041 and 0.174 for accuracy, precision, recall, specificity and F1-score respectively. As mentioned above, the multi-class accuracy as well as the specificity are misleading metric and the differences obtained in precision, recall and F1-score were expressive.

This indicates that in this experiment TERL was not capable to generalize during training to correctly classify the sequences from different databases during test and surpass the PASTEC results. These results are possibly due to the occurrence of wrongly annotated sequences and sequences with long unidentified inter-contig regions as noise in the test set.

However, it is important to highlight that TERL used just a fraction of the RepBase to train and TEclass and PASTEC use the entire database as homology search source. This could lead to an advantage for these methods.

Table 13 presents a summary of the results obtained in the experiments 2, 4 and 5. All metrics are the macro mean obtained by the methods. This table allows a comparison between the results of all analyzed methods. In this table we can also see the difference about the test time between the methods and how fast TERL is since it classified the sequences in about 1.10 seconds against 1.65 × 10^4^ and 1.66 × 10^2^ seconds for PASTEC and TEclass, respectively. TERL is four orders of magnitude faster than PASTEC and about 150 times faster than TEclass.

**Table 13.**
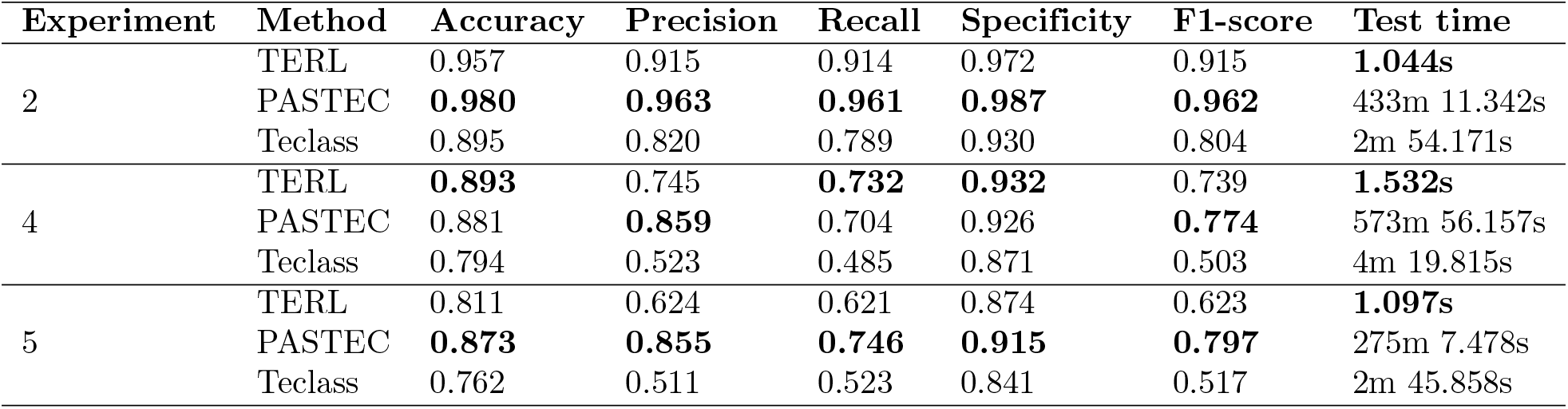
Overall performance metrics obtained by each method for experiments 2, 4 and 5. All values are the macro mean of the respective metric.

| Experiment | Method | Accuracy | Precision | Recall | Specificity | F1-score | Test time |
| --- | --- | --- | --- | --- | --- | --- | --- |
| 2 | TERL | 0.957 | 0.915 | 0.914 | 0.972 | 0.915 | **1.044s** |
|  | PASTEC | **0.980** | **0.963** | **0.961** | **0.987** | **0.962** | 433m 11.342s |
|  | TEclass | 0.895 | 0.820 | 0.789 | 0.930 | 0.804 | 2m 54.171s |
| 4 | TERL | **0.893** | 0.745 | **0.732** | **0.932** | 0.739 | **1.532s** |
|  | PASTEC | 0.881 | **0.859** | 0.704 | 0.926 | **0.774** | 573m 56.157s |
|  | TEclass | 0.794 | 0.523 | 0.485 | 0.871 | 0.503 | 4m 19.815s |
| 5 | TERL | 0.811 | 0.624 | 0.621 | 0.874 | 0.623 | **1.097s** |
|  | PASTEC | **0.873** | **0.855** | **0.746** | **0.915** | **0.797** | 275m 7.478s |
|  | TEclass | 0.762 | 0.511 | 0.523 | 0.841 | 0.517 | 2m 45.858s |

The difference showed in the results obtained by all methods between the classification of experiment 2 and 5 indicates that the RepBase database does not provide enough patterns to allow machine learning algorithms to achieve good generalization.

Part of this can be due the RepBase sequences rarely presenting unidentified intercontig regions and other databases presenting sequences from different organisms, which could negatively affect the performance of the methods, due to the use of homology searches as feature.

Experiment 6 verifies the performance of the PASTEC method when some of the homology based search files are not provided. Table 14 shows the metrics obtained by the PASTEC on the classification of data sets 2, 4 and 5.

**Table 14.**
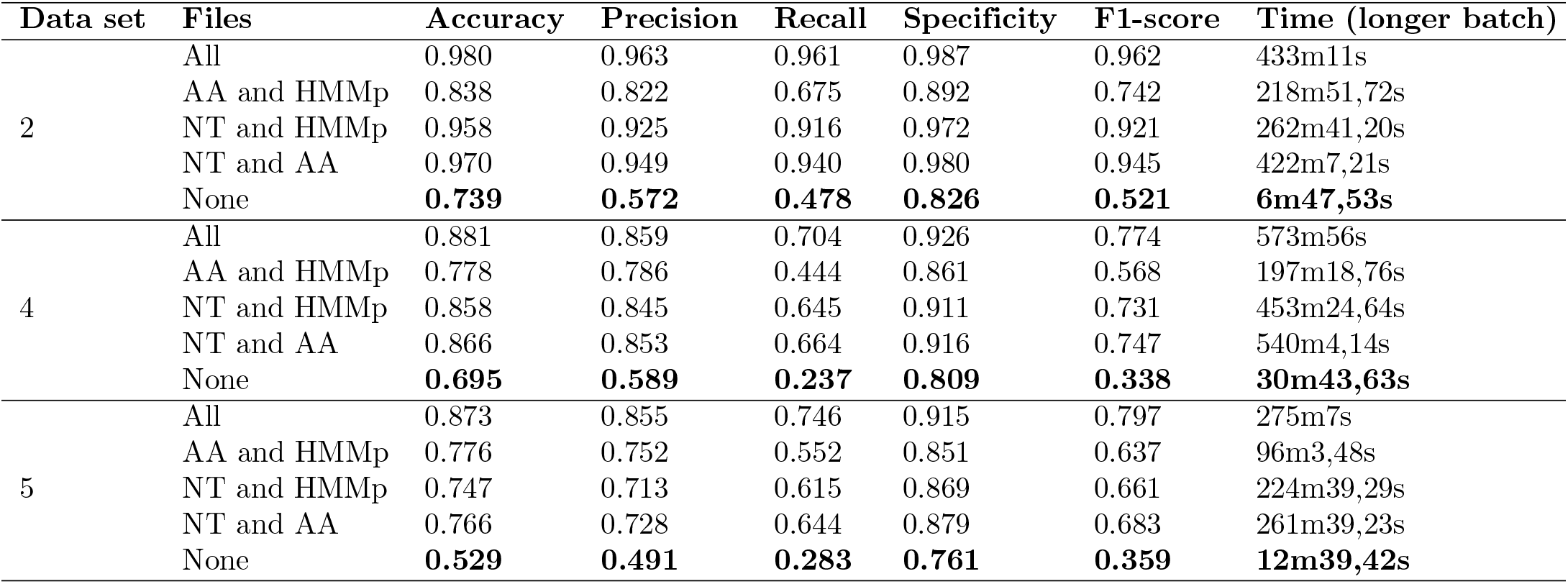
Ablation of PASTEC by removing auxiliary homology based search files on experiment 6. The auxiliary homology based search files are: RepBase nucleic acid sequences (NT), RepBase amino acid sequences (AA) and HMMs profiles (HMMp) from Gypsy Database 2.0. This results shows that the PASTEC method is strongly dependent on these auxiliary files.

| Data set | Files | Accuracy | Precision | Recall | Specificity | F1-score | Time (longer batch) |
| --- | --- | --- | --- | --- | --- | --- | --- |
| 2 | All | 0.980 | 0.963 | 0.961 | 0.987 | 0.962 | 433m11s |
|  | AA and HMMp | 0.838 | 0.822 | 0.675 | 0.892 | 0.742 | 218m51,72s |
|  | NT and HMMp | 0.958 | 0.925 | 0.916 | 0.972 | 0.921 | 262m41,20s |
|  | NT and AA | 0.970 | 0.949 | 0.940 | 0.980 | 0.945 | 422m7,21s |
|  | None | **0.739** | **0.572** | **0.478** | **0.826** | **0.521** | **6m47,53s** |
| 4 | All | 0.881 | 0.859 | 0.704 | 0.926 | 0.774 | 573m56s |
|  | AA and HMMp | 0.778 | 0.786 | 0.444 | 0.861 | 0.568 | 197m18,76s |
|  | NT and HMMp | 0.858 | 0.845 | 0.645 | 0.911 | 0.731 | 453m24,64s |
|  | NT and AA | 0.866 | 0.853 | 0.664 | 0.916 | 0.747 | 540m4,14s |
|  | None | **0.695** | **0.589** | **0.237** | **0.809** | **0.338** | **30m43,63s** |
| 5 | All | 0.873 | 0.855 | 0.746 | 0.915 | 0.797 | 275m7s |
|  | AA and HMMp | 0.776 | 0.752 | 0.552 | 0.851 | 0.637 | 96m3,48s |
|  | NT and HMMp | 0.747 | 0.713 | 0.615 | 0.869 | 0.661 | 224m39,29s |
|  | NT and AA | 0.766 | 0.728 | 0.644 | 0.879 | 0.683 | 261m39,23s |
|  | None | **0.529** | **0.491** | **0.283** | **0.761** | **0.359** | **12m39,42s** |

For the experiment 6 (Table 14), we executed for data sets 2, 4 and 5 a series of tests removing each time one of the auxiliary files and as the final test we removed all files. For each data set test, first we show the results obtained by using all homology searches, on the second test we removed the RepBase nucleotide source file (NT), on the third test we removed the amino acid source file (AA), on the fourth test we removed the HMMs profiles file (HMMp) and on the last test we remove all files.

The obtained results show the dependancy of these files as the metrics begin to drop as files are removed. We can also observe which file apparently was providing more evidences for the classification. Except for the test with data set 5, the files that show to improve the results are the nucleotide and amino acid source files. The HMMs profiles source only helped improve the accuracy and precision on the test with data set 5.

The homology search in the nucleotide and amino acid source files also showed to be the more time consuming steps of the PASTEC classification.

Comparing the results with no homology search, all PASTEC results are worse than the ones obtained by TERL. Also all times are worse than the ones obtained by TERL, showing that TERL does not need any additional prior information, just the sequences, and can achieve much better results in some cases, if the use of homology search is considered, and in all cases if no homology search is consider.

## Conclusions

This paper presents a pipeline that can classify transposable elements in all levels of hierarchy and other biological sequences.

Different from other approaches, the proposed method does not rely on homology search and other manually-defined features, which affects negatively the performance when classifying non-homologous sequences or sequences with different assembly qualities (as shown in the results of experiment 5 on Figure 8 and Table 12), thus providing a general method that can classify any type of sequence.

As verified, different databases can present duplicated sequences and same sequences classified as different classes, which could negatively affect machine learning algorithms. Thus a preprocessing step is needed to address these problems. We present some preprocessing steps that can be applied to address these problems and prepare DNA sequences to classification by a deep CNN, which is the two-dimensional space transformation (i.e., step 5 in Figure 1).

The results of the experiments 1 and 2 show that the proposed pipeline for TERL is able to classify well-annotated and curated TE sequences into superfamily and order levels and present very high accuracy and specificity. The precision, recall and F1-score obtained also indicate that TERL is a good alternative to classify TE sequences.

Experiments 3 and 4 indicate that adding sequences from other databases in the train and test sets present a more challenging classification problem since the metrics obtained by all methods dropped. This can be due to the insertion of sequences from different organisms, sequences with different sequencing qualities (i.e. sequences that can present unidentified inter-contig regions) and sequence with the wrong annotation. Even so, the proposed method obtained better metrics in the classification on experiment 4.

Experiment 5 shows that RepBase sequences do not present enough patterns to train the CNN used on TERL to the point that it generalizes and surpasses the metrics obtained by the other methods. This can be due to the fact that RepBase sequences rarely present long unidentified regions of *N* symbols, which often occurs in the sequences from the other databases.

PASTEC method showed to be very time consuming and depends on auxiliary homology search files. These homology searches are responsible for great part of the computation time of PASTEC.

Deep CNNs showed to far surpass all methods upon test time since the test times of the entire test sets were less than 2 seconds. This is due to GPU acceleration provided by frameworks such as Tensorflow.

For future works, we intend to study the performance of recurrent neural networks, such as LSTM, to classify biological sequences, given it is a well known sequence classifier [13] [20] [22].

## Acknowledgments

We gratefully acknowledge the support of NVIDIA Corporation with the donation of the Titan V GPU used for this research.

